# Joint analysis of human retroelements-linked histone modification profiles reveals quickly evolving molecular processes connected with cancer

**DOI:** 10.1101/2025.09.24.677146

**Authors:** Daniil Nikitin

## Abstract

Human retroelements (REs), which comprise approximately 40% of the genome, have played a pivotal role in the evolution of key molecular processes, such as placental development, by introducing novel regulatory elements near host gene promoters and enhancers. Despite their genomic abundance and regulatory influence, the functional trajectories of REs remain poorly understood. Here, leveraging ChIP-seq profiles of histone modifications (H3K4me1, H3K4me3, H3K9ac, H3K27ac, H3K27me3, and H3K9me3) from five human cell lines deposited in the ENCODE database, we systematically ranked the regulatory impact of REs across 25,075 human genes. Gene sets enriched for promoter- and enhancer-associated RE-linked regulatory sites were identified. Consensus gene sets across cell lines were found to be associated with pathways involved in cancer progression, specifically chronic myeloid leukemia and small cell lung cancer, as well as with host defense responses to infection with human T-cell lymphotropic virus type 1. These findings provide new insights into recent human evolution and highlight the ongoing influence of selfish genetic elements on genome regulation and disease susceptibility.

## Introduction

Human retroelements (REs, also called retrotransposons), namely LTRs, SINEs and LINEs, comprise 40% of the genome [1], [2]. They impact the host genome by providing regulatory sites [3] and novel protein coding genes, manifesting themselves in various human diseases [4]. For example, human endogenous retroviruses (LTR) derived protein coding genes suppress maternal anti fetal immune response in placenta (like syncytin-1 [5]). All REs transpose via an RNA intermediate and then a DNA copy is synthesised and inserted into other genomic locus. Therefore, REs bear transcription factor binding sites enriched with active chromatin marks (H3K4me1, H3K4me3, H3K29ac etc) that can rewire the host gene regulatory networks [6]. On the other hand, REs can be repressed by the host defense mechanisms, such as CpG methylation, which allows to mitigate deleterious effects of new genetic elements insertion near essential genes [7]. In a greater perspective, there is an evolutionary arms race between mobile selfish genetic elements and host defensive systems [8], which leads to such complex innovations as adaptive immunity, including both the microbial (CRISPR-Cas) and the vertebrates one (VDJ-recombination) [8].

In order to decipher the evolutionary impact of REs on human gene regulation, in 2018-2019 we analyzed genome profiles of transcription factors and chromatin modifications in connection with RE loci and their impact onto the adjacent gene expression. In 2018 we studied 225 transcription factors binding profiles in the K562 cell line and measured regulatory impact of human REs, including the evolutionary young ones (inserted after the radiation of Old World monkeys) [1]. We found that such molecular processes as PDGF, TGF beta, EGFR, and p38 signaling were positively regulated by evolutionary young REs [1]. In 2019 we expanded the analysis, profiling 563 transcription factors binding in 13 cell lines [9], as well as the enhancer-specific chromatin modification H3K4me1 [10] and simultaneously five chromatin marks (H3K4me3, H3K9ac, H3K27ac, H3K27me3 and H3K9me3 [11]). We developed the original method called RetroSpect and showed that such human molecular processes as gene regulation by microRNAs, olfaction, color vision, fertilization, cellular immune response, and amino acids and fatty acids metabolism and detoxification were enriched with RE-specific regulation and hence were quickly evolving under the RE pressure [9].

Despite the significant progress being made, the field remains generally unexplored. Chromatin marks form a complex interactive system with readers, writers and erasers, collectively regulating gene expression [12]. In the previous articles we gathered gene-level RE regulatory impact scores for the following chromatin marks:

- H3K4me1 - active enhancer specific [13]
- H3K27ac - active enhancer specific [14]
- H3K4me3 - promoter specific [15]
- H3K9ac - promoter specific [16]
- H3K27me3 - polycomb repression specific [17]
- H3K9me3 - heterochromatin specific [18]

In the present work we jointly analyse active promoters, enhancers and heterochromatin RE-linked chromatin marks at the level of genes, based on the fact that REs can both activate and inactivate adjacent human genes because the defensive host heterochromatinisation which is relevant in pathology, for example in X-linked dystonia parkinsonism [19]. The proposed comprehensive intersection analysis based on the original RetroSpect method allowed to show that RE-linked promoter histone marks activate chronic myeloid leukemia and small cell lung cancer connected genes in all 5 cell lines under study. Moreover, the RE-linked promoter regulation impacts such general cancer associated processes as cell cycle and cellular senescence. Surprisingly, we observed RE-linked upregulation of human T-cell leukemia virus type 1 infection related genes and the level of promoter specific histone modifications. In contrast, active RE-linked enhancers are more cell type specific and do not form cell line consensus groups of enriched processes. Taken together, these findings push forward our understanding of human RE-impacted evolution of gene regulatory networks in health and disease.

## Materials and methods

### Gene-level scores

Scores of RE-linked epigenetic marks regulatory impact were downloaded for 6 chromatin modifications (H3K4me1, H3K27ac, H3K4me3, H3K9ac, H3K27me3 and H3K9me3) and for 5 cell lines (K562, HepG2, GM12878, MCF-7, HeLa-S3) from the previously published papers [10], [11]. The scores were calculated based on more than 1.5 million histone tags for 25075 genes. The following scores were used in the current research:

- GRE, gene RE-linked enrichment score.
- NGRE, normalized gene RE-linked enrichment score.

Formulas and biological meaning of these scores are described in Nikitin et al., 2019a [10].

### Selection of RE-linked regulatory impact enriched and deficient genes

According to the original RetroSpect approach [9], for each chromatin modification in each cell line we plotted NGRE vs GRE in a scatter plot, build a trend line by the method of least squares [20], and top-10% and bottom-10% genes by their distance to the trend line were selected. Genes from the top-10% group (with NGRE score high compared to the GRE one) were considered RE-enriched, the latter group was deemed RE-deficient.

### Plots and visualizations

Plots were drawn using the python matplotlib [21] and seaborn [22] packages, Venn diagrams were drawn using the python supervenn [23] package. Principal component analysis (PCA) with centroid approximation was done using the python scikit-learn package [24].

### Gene Ontology analysis

Gene Ontology analysis was done using the ShinyGo service from South Dakota State University [25].

## Results and discussion

### Dimensional reduction and major data properties

In order to study the degree of cell-specific and chromatin modification-specific differences for the dataset of NGRE and GRE scores in 6 chromatin modifications and 5 cell lines, we performed principal component analysis with centroid approximation (Fig. 1).

**Figure 1.**
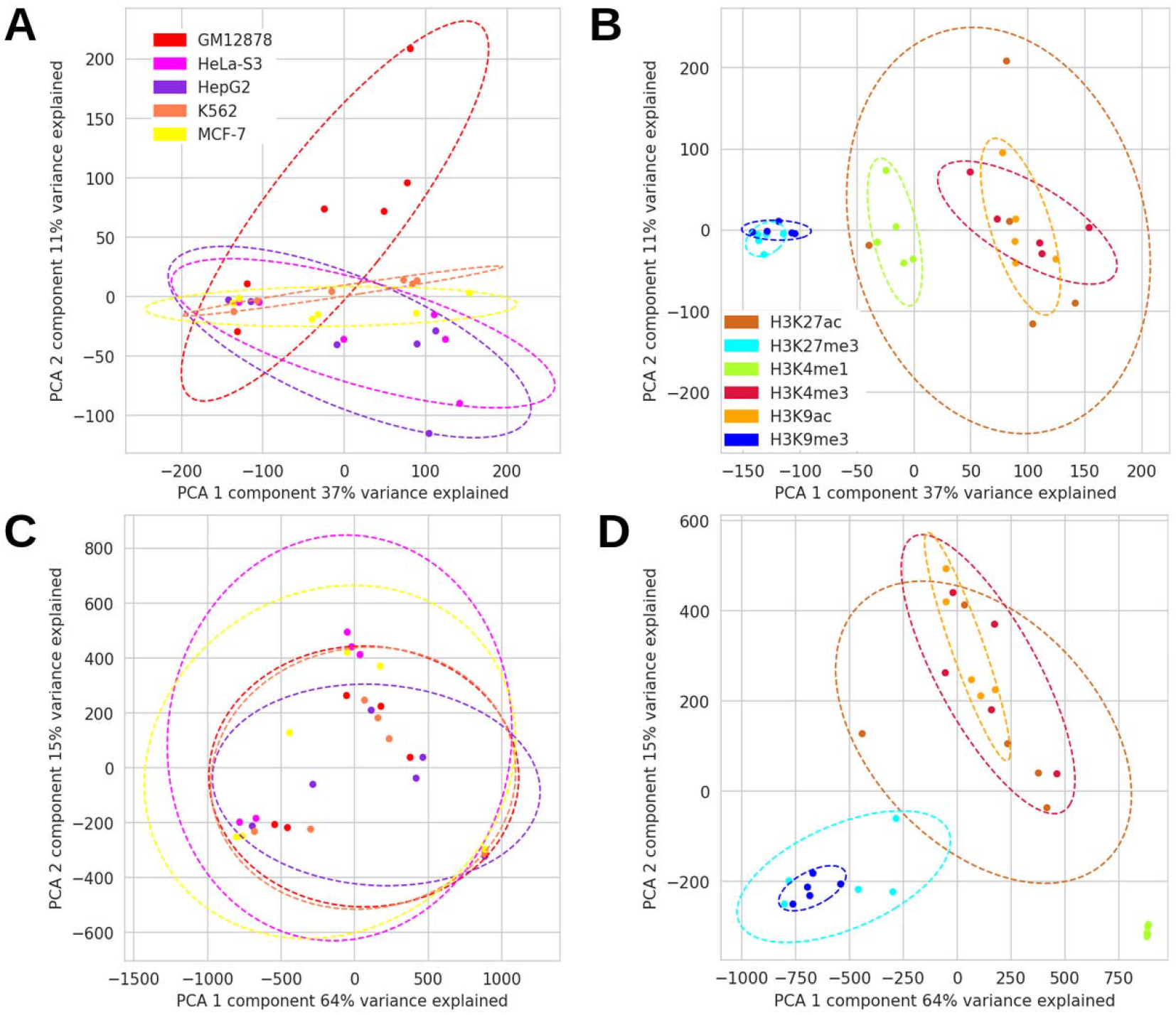
Principal component analysis of GRE and NGRE scores. A - PCA of GRE colored by cell line, B - PCA of GRE colored by histone modification, C - PCA of NGRE colored by cell line, D - PCA of NGRE colored by histone modification. Centroid approximations are drawn with 95% confidence intervals. The palettes for cell lines in A and C are the same, as well as the palettes for chromatin modifications in B and D.

Mapping of different cell lines with the same histone modifications intersects to a high degree (Fig. 1B, 1D) with heterochromatin marks clustering differently compared to the active chromatin ones both for GRE (Fig. 1B) and NGRE (Fig. 1D). Notably, H3K4me1, the active enhancer mark, is mapped separately in both cases, and the second active enhancer one, H3K27ac, is located the most highly differing between the cell lines. Contrastingly, different cell lines do not form isolated groups, albeit they are from different organs and tissues (Fig. 1A, 1C). This corresponds with fundamental differences between chromatin marks in terms of their functioning, whereas cell line specific patterns are less pronounced.

### Selection of RE-enriched and deficient genes for individual cell lines and chromatin modifications

According to the original RetroSpect procedure [10], for all 5 cell lines and 6 chromatin modifications we correlated NGRE and selected RE-enriched and deficient genes (Fig. 2).

**Figure 2.**
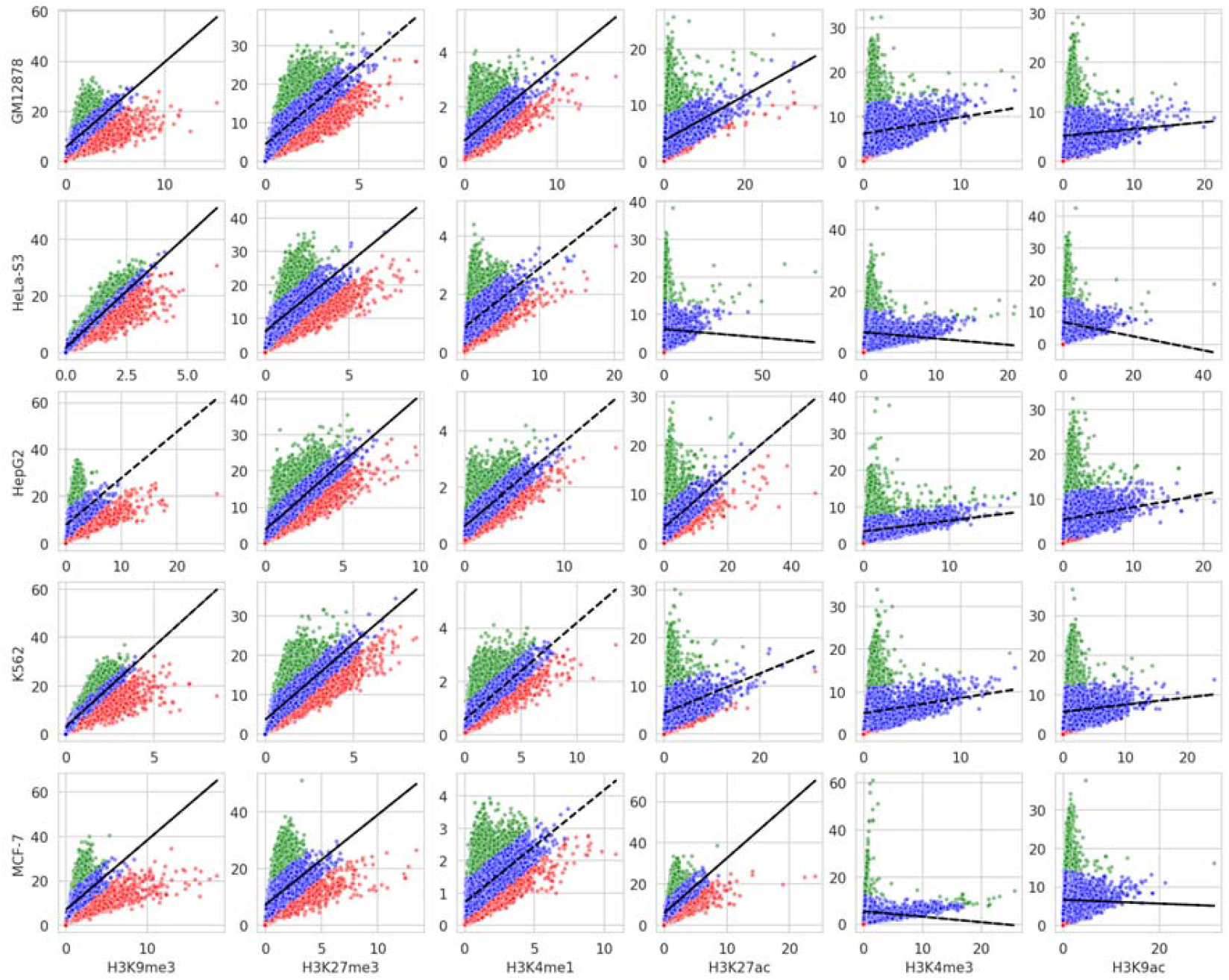
Correlations of NGRE (y-axis) and GRE scores for 5 cell lines (shown by rows, names written on the leftmost part of the plot grid) and 6 chromatin modifications (shown by rows, names written on the bottom part of the plot grid). For each plot the linear trend line was built using the least squares method, and top 10% genes with the highest distance to the curve from above are shown in green (RE-enriched genes), top 10% genes with the highest distance to the curve from below are shown in red (RE-deficient genes) and the rest 80% genes are blue.

Out of 30 all combinations, 5 ones showed negative correlations because of two highly divergent linear trends: H3K27ac, H3K4me3, H3K9ac in the cell line HeLa-S3, and H3K4me3, H3K9ac in the cell line MCF-7. The two heterochromatin histone modifications (H3K27me3 and H3K9me3) and the enhancer specific one H3K4me1 showed no negative linear trends, mimicking the patterns observed earlier by us [9]. The major difference of the current approach applied here is the fact that RE-enriched and deficient genes are extracted individually by cell lines and chromatin modification, whereas earlier the GRE scores of different cell lines and epigenomic features (such as transcription factors) were averaged [10]. At this stage of the analysis the functional significance of the selected RE-enriched and deficient genes is unclear, so further intersections and Gene Ontology approaches were applied as described below.

### Intersection of the RE-enriched and deficient genes based on different cell line and histone modifications

In order to understand how REs impact human gene regulation via different histone modifications and in different cell lines, we intersected RE-enriched and deficient genes by different modifications with each other, repeating the procedure in all 5 cell lines (Fig. 3).

**Figure 3.**
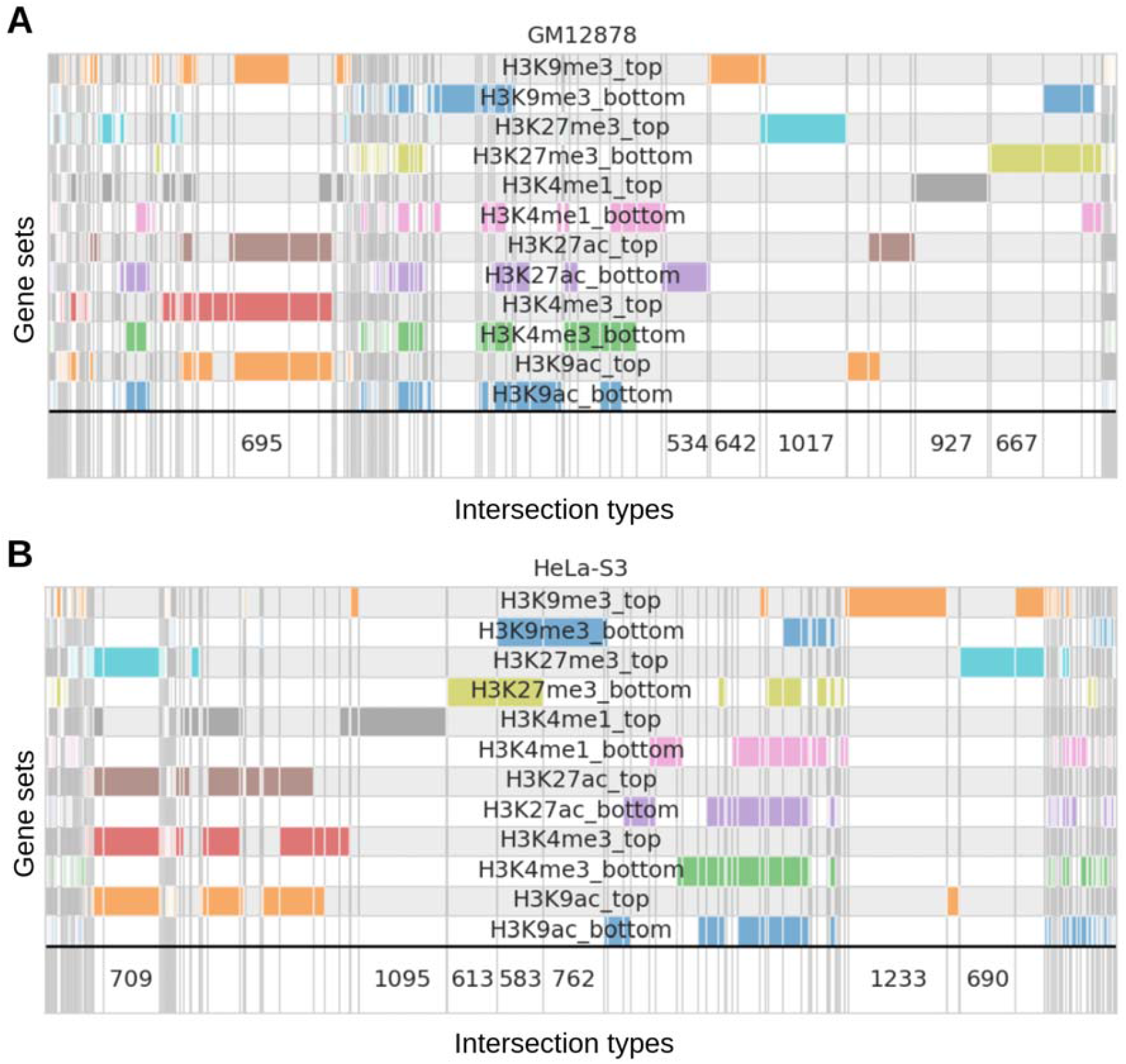
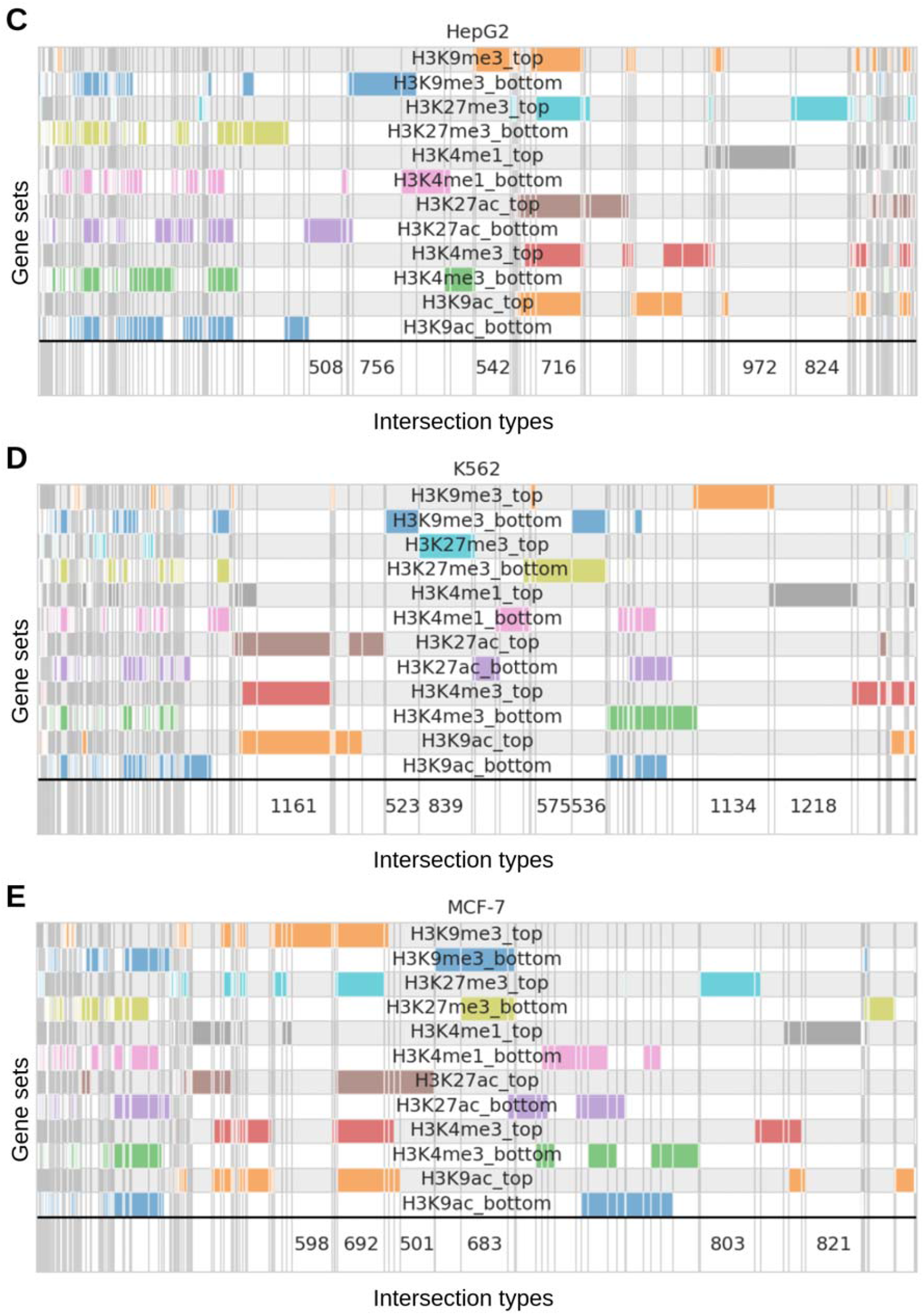
Supervenn plot showing intersections between RE-enriched (abbreviated as ‘top’) and RE-deficient (abbreviated as ‘bottom’) for each histone modification (shown on each row) for each cell line. A - GM12878, B - HeLa-S3, C - HepG2, D - K562, E - MCF-7. Each gene set is shown using a different color. Colored grey rectangles denote the successful intersection (common genes), white and light grey ones indicate no intersection.Intersection groups that have more than 500 genes have gene numbers written in the bottom of each panel.

The intersection analysis shows that different cell lines have different interplay of RE-linked active and repressive chromatin marks. For example, the K562 cell line (Fig. 3D) has a large cluster of genes (1161) that share promoter and enhancer active marks (H3K27ac, H3K4me3, H3K9ac) without any heterochromatin marks - this can be interpreted as genes that are actively regulated by REs. In contrast, in MCF-7 (Fig. 3E) such a cluster is less than 500 genes, and 692 genes are RE-enriched by active marks (H3K27ac, H3K4me3, H3K9ac) and both repressive marks simultaneously, which pinpoints ambiguity of RE-linked epigenetic regulation of the host genes. The same situation is observed in HepG2 (716 genes, Fig. 3C), whereas in GM12878 (Fig. 3A) and HeLa-S3 (Fig. 3B) both types of clusters are present: the one with the active marks only and the one with active and repressive marks (either H3K9me3 or H3K27me3, respectively). This can be connected with the fact the RE evolutionary pressure is exerted at different intensities between tissues, with placenta [26] and neocortex [27] being one of the most invaded by REs (transcriptionally active) and hence quickly evolving in the human lineage.

### Promoter and enhancer-level RE-enriched genes selection among the five cell lines

Having the dataset of GRE and NGRE for 5 cell lines and 6 histone modifications, we got 12 sets of RE-enriched and deficient genes in each cell line. In order to get cell line consensus of promoters and enhancers specific RE-enriched genes, we defined enhancers and promoters RE-enriched genes in the following way for each cell line:

- Enhancer specific genes that are intersection of RE-enriched genes by H3K4me1 and H3K27ac, and RE-enriched genes by heterochromatin marks (H3K9me13 and H3K27me3) are subtracted from this intersection.
- Promoter specific genes that are intersection of RE-enriched genes by H3K4me3 and H3K9ac, and RE-enriched genes by heterochromatin marks (H3K9me13 and H3K27me3) are subtracted from this intersection in the same way.

For each cell line we obtained a variable number of promoter and enhancer RE-enriched genes, and we intersected them between all the cell lines to get the degree of cell-specificity in RE-linked genes regulation and regulatory evolution (Fig. 4).

**Figure 4.**
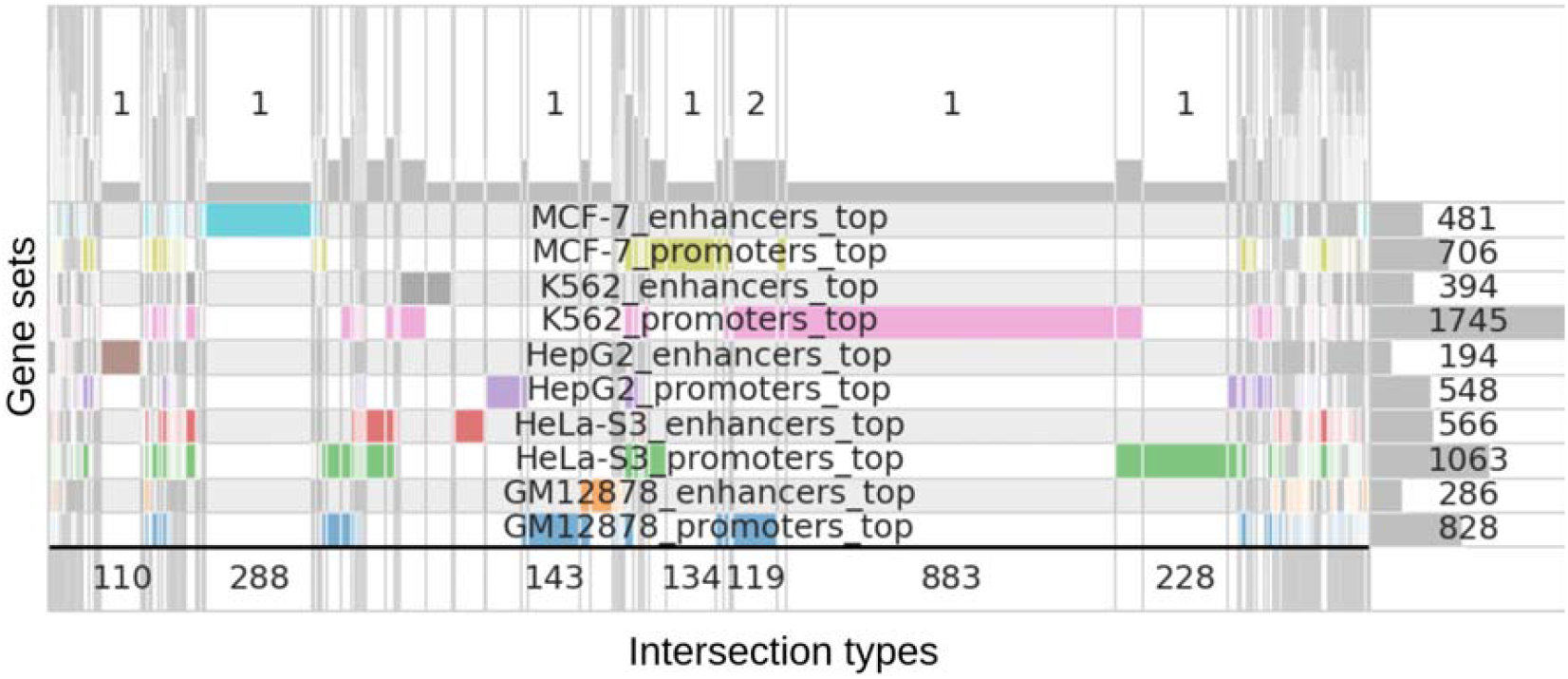
Supervenn plot showing intersections of enhancer (enhancers_top) and promoter (promoter_top) specific RE-enriched genes in the 5 cell lines investigated in the current study. The numbers on the right part of the plot are indicating gene numbers in the sets shown. Each gene set is shown using a different color. Colored grey rectangles denote the successful intersection (common genes), white and light grey ones indicate no intersection. Intersection groups that have more than 100 genes have gene numbers written in the bottom of each panel. Numbers on the top part of the plot are showing the number of gene sets that share the intersection (for cases with more than 100 genes).

The intersection patterns shown in Fig. 5 illustrate the degree of concordance between RE-enriched promoter- and enhancer-associated regulation across different cell lines. Notably, 92 promoter-associated RE-enriched genes were shared across all five cell lines (ranging from 548 to 1,745 genes per cell line), whereas only three enhancer-associated RE-enriched genes were common (from 194 to 566 genes per cell line). This disparity likely reflects the greater variability of enhancer-associated chromatin modifications (H3K4me1 and H3K27ac; Fig. 3), as well as the well-established observation that transcriptional regulation at promoters is more conserved across tissues than at enhancers [28]. Furthermore, promoter- and enhancer-specific RE-enriched gene sets exhibited minimal overlap within each cell line, suggesting that distinct subsets of REs - potentially corresponding to different RE classes - modulate gene expression at the promoter and enhancer levels, respectively. This hypothesis warrants further investigation through integrative computational analyses and experimental validation.

### Functional characterisation of the cell line consensus promoter and enhancer-level RE-enriched genes

The consensus RE-enriched promoter and enhancer genes are shown in the Table 1 below.

**Table 1.**
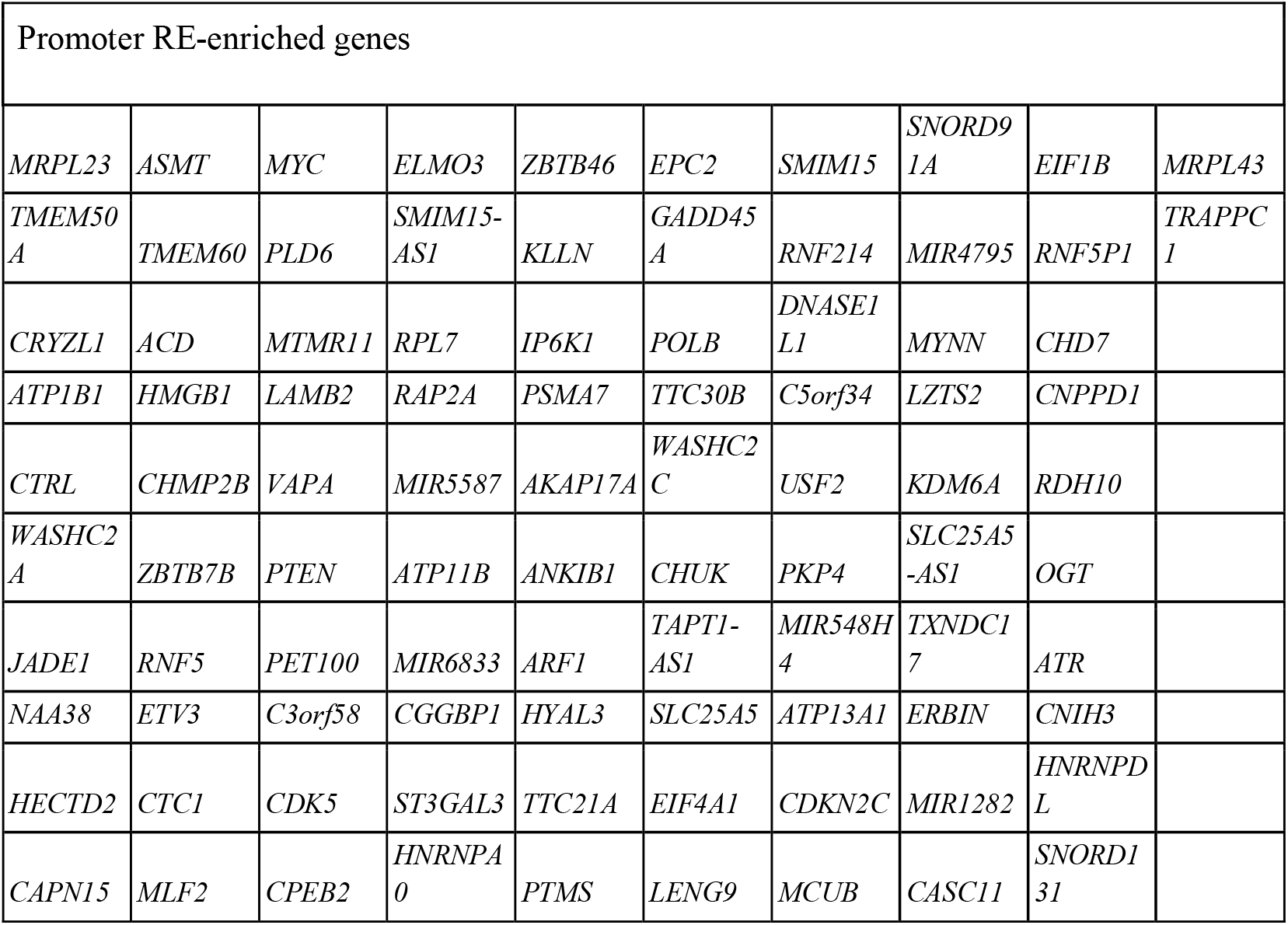
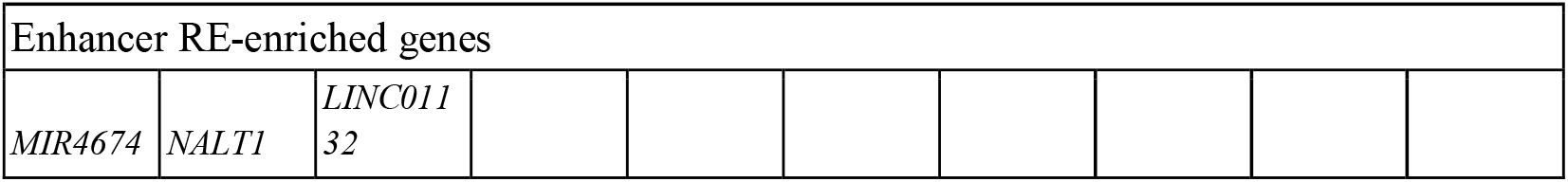
The list of consensus RE-enriched promoter and enhancer genes.

The genes found were tested for functional enrichment using the Gene Ontology approach [29] via the ShinyGO portal [25] without the background genes, and the enrichment processes were filtered by False Discovery Rate [30] threshold 0.1. The resulting processes for the RE-enriched promoter genes are shown in Table 2.

**Table 2.** The list of enrichment results for RE-enriched promoter genes.

| Enrichment FDR-corrected p-value | Number of genes from RE-enriched promoters in the process | Number of genes in the process | Fold Enrichment | The process with the GO knowledgebase hyperlink |
| --- | --- | --- | --- | --- |
| 0.02 | 4 | 92 | 12.6 | Small cell lung cancer |
| 0.064 | 3 | 76 | 11.4 | Chronic myeloid leukemia |
| 0.044 | 4 | 126 | 9.2 | Cell cycle |
| 0.016 | 6 | 222 | 7.8 | Human T-cell leukemia virus 1 infection |
| 0.064 | 4 | 156 | 7.4 | Cellular senescence |

Gene Ontology analysis (Table 2) revealed that human REs transcriptionally activate genes associated with two major cancer types - small cell lung cancer and chronic myeloid leukemia - as well as genes upregulated during infection with human T-cell leukemia virus type 1 (HTLV-1), a retrovirus that, similar to HIV, targets CD4□ T cells [31]. These findings raise the possibility that RE-mediated regulatory activity and evolutionary selection may operate, at least in part, through the transcriptional repurposing of pre-existing host defense mechanisms, a phenomenon exemplified by placental evolution [5]. Additional enriched gene sets, including cell cycle regulation and cellular senescence - both canonical hallmarks of cancer [32] - further support the notion that promoter-associated, RE-enriched regulatory programs are linked to oncogenesis. By contrast, no statistically significant functional enrichments were observed for the three enhancer-associated, RE-linked genes identified (*MIR4674, NALT1*, and *LINC01132*), two of which are non-coding.

## Conclusion

We investigated the regulatory impact of retroelements (REs) on human gene regulation by analyzing promoter-associated, enhancer-associated, and heterochromatin-associated histone modifications, using ChIP-seq whole-genome profiles from five human cell lines. Clustering and intersection analyses revealed that RE-associated regulatory activity at enhancers exhibits a high degree of cell type specificity, providing a foundation for further studies on the mechanisms underlying this specificity. In contrast, RE-associated regulation at active promoters appeared to be more conserved across different cell types. Notably, REs were found to activate pathways related to cancer-associated processes, including cellular senescence and cell cycle regulation. These pathways are linked to cancer diseases such as small cell lung cancer and chronic myeloid leukemia, and RE-associated regulatory activity also impacts host cell defense mechanisms against human T-cell lymphotropic virus type 1. Collectively, these findings advance our understanding of the role of REs in human evolution and oncogenesis.

